# Optimizing reduction of Western blotting analytical variations: use of replicate test samples, multiple normalization methods, and sample loading positions

**DOI:** 10.1101/2020.12.16.423026

**Authors:** Phyllis A. Rees, R. Joel Lowy

## Abstract

Western blot (WB) analysis is widely used, but obtaining consistent results can be problematic, particularly if all the treatments cannot be placed on one gel. Sources of variation were investigated by analyzing a set of samples which should produce statistically identical results, an approach commonly applied to analytical instruments. Test samples were lysates from RAW 264.7 murine macrophages treated with LPS to activate MAPK and NF-kB signaling targets. The resulting WB were probed for p-ERK, ERK, IkBβ and non-target protein levels. The first test set (Set 1) was a pooled cell lysate and the second (Set 2) four independently prepared lysates.

Analytical replicate data were generated by using multiple WB. Different normalization methods and sample groupings were applied to the density values and the resulting coefficients of variation (CV) and ratios of maximal to minimal values (Max/Min) were compared. Ideally, as the samples were identical (Set 1), or nearly so (Set 2), CVs would be 0 and the Max/Min 1. Results showed the raw density data variance was composed of variability between gel lanes and between gels; but only between gels after applying normalization calculations. CV and Max/Min were 30-70% and 1.5 for raw data, respectively. Common normalization methods, total lane protein, % Control, and p-ERK/ERK ratios, did not have the lowest CVs or Max/Min values. Some of these methods were counterproductive, greatly increasing CV and Max/Min. The best normalization methods resulted in CV and Max/Min value, as low as 5-10% and 1.1, respectively, but more typically 20-25% and 1.2. Application of these methods should allow reliable interpretation of complex experiments that require samples to be placed on multiple gels to provide results, even if fold changes in the target proteins are relatively modest.

## Introduction

Western blotting (WB) is primarily a binary or very broad semi-quantitative comparison method for demonstrating a large change in a protein of interest. Analytical, also termed technical, variation introduced by the method itself generally does not interfere with obtaining useful results. However, cell and molecular biology continue to ask increasingly more complex questions that require target proteins to be determined simultaneously for multiple conditions in parallel, with increasing accuracy, and for smaller proportional changes. These studies also often require semi-quantitative or full quantitative analysis of the actual amount of protein present. All of these demands place more stringent requirements on the analytical methodological variation that can be tolerated from WB [1, 2]. A number of reviews have thoroughly examined the various steps in the process and identified improvements or requirements for procedures to obtain results which are more reproducible between different sets of analyses within studies and between different studies [1, 3-9]. Particular emphasis has been placed on the linearity of detectors [4, 10, 11], antibody characterization and concentrations [2, 5, 12, 13], background subtraction [1, 11], standard curves for quantitation [14-17], and between-lane normalization methods [18]. A smaller group of studies have addressed post acquisition methods of densitometry analysis, [15, 19] and/or quantification of errors [20-23], and improvement of time series data [21] to reduce WB data variance, but do not seem to have been widely adopted.

In this study the method commonly used to test analytical instruments is applied to the WB process. In general to verify proper function of instruments, samples of identical known composition and/or known concentration are repetitively measured. Standards with a previously established value allow the absolute accuracy of the instrument to be determined. Replicate measurement of identical samples, often referred to as analytical or technical replicates, establishes reproducibility, e.g. precision. Often the approaches are combined by replicate analysis of an established standard. Standard and analytical replicates samples are also used for instruments with parallel channels, such as multi-well plate spectrophotometers, to verify uniformity of responses within the specified capabilities of the instrument. All these types of analyses are used to make statements about the sensitivity and reliability of the instrument’s measurement capability and expressed as mean values and ranges of deviation.

In this study to test the reproducibility of WB all lanes of replicate electrophoresis gels were loaded with the same pooled sample and levels of target protein determined by standard WB methods. In a second set of assays samples from replicate independent experiments were used to add an experimental variability component, but again were loaded in replicate lanes on replicate gels. Using these data several different methods of data normalization were compared for their efficacy in controlling variation of the WB densitometry results. In addition, the experimental design allowed testing of whether different patterns of sample loading could contribute to controlling WB analytical variations.

## Methods

### Cell Culture and Reagents

RAW 264.7 cells (RAW, ATCC, Manassas, VA) were propagated in Dulbecco’s modified Eagles medium with 4.5 mg/ml glucose (DMEM, #10313-021, Invitrogen, ThermoFisher, Pittsburgh, PA), supplemented with 10% heat inactivated fetal bovine serum (FBS, HyClone, GE Healthcare, South Logan, UT #SH30070.03), 2 mM L-glutamine (Invitrogen, #25030-081) and 100 U/ml Penicillin and 100 ug/ml Streptomycin (P-S, Invitrogen, 15140-122). LPS, *E. coli* serotype 0127:B8, (Sigma, St. Louis, MO #L-4516) was used at a final concentration of 1 ug/ml in all experiments.

### Preparation of Lysates and Western Blotting

RAW 264.7 cells were plated at a density of 1 x 10^6^ cells/ml, 2 ml per well in a 6 well plate. Cells were plated 16 – 24 hours prior to the experiment to allow settling. LPS or control treatment was done by first removing old media and then adding 2 ml media + 1 ug/ml LPS per well to 3 wells or adding media only to 3 wells. Incubation time was 1 hour in a 37°C, CO_2_ incubator, after which media was removed and the cell monolayer washed 1 time with 2 ml cold DPBS with calcium and magnesium (DPBS++, Invitrogen, #14040-133). The DPBS++ was removed and 100 ul of ice cold lysis buffer was added containing: 100 mM Tris-HCL (Quality Biological Inc., Gaithersburg, MD, #351-007-101) pH8.0 at 25°C; 100 mM NaCl, (Sigma #S3014); 2 mM EDTA, (Sigma, #E5134); 1% v/v 10% Igepal CA (Sigma, #CA630); 1 mM sodium vanadate (New England Biolabs, Ipswich, MA, #0758s); 50 mM sodium fluoride (Sigma, #S7920); 1X cOmplete mini EDTA free protease inhibitor solution (Roche, Branford, CT, #11836170001). The lysates were harvested and immediately placed on ice. Lysates were centrifuged for 10 minutes at 16,168xg, 4°C to remove cellular debris, transferred to fresh tubes and were immediately frozen at -80°C. Prior to electrophoresis, the protein concentration was determined using the BioRad DC microplate assay protocol (BioRad DC Protein Assay Kit, BioRad, Hercules, California #5000112). Protein Standards and samples were run on NuPAGE 10% Bis-Tris Gels (Invitrogen, #NP0315) for 2 hours at 100V. Two protein standards were run in the first two lanes of each gel: BioRad Kaleidoscope Pre-stained Standard (BioRad, #161-0324) and Magic Mark XP Western Standard (Invitrogen, #LC5602). Lysates were loaded with control (LPS untreated) and LPS treated samples alternately in the next 8 lanes (see S3 Figure). After electrophoresis the proteins were transferred to Nitrocellulose membranes (Amersham Protran Premium Nitrocellulose, GE Healthcare, #10600048) via wet transfer at a current of 45V for approximately 20 hours at 4°C. After transfer the gels were stained with Gel Code Blue (Thermofisher, #24952) for 1 hour and then destained with water for 1 hour. The nitrocellulose membranes, after transfer, were fixed in isopropanol for 1 minute. Membranes were quickly washed 3 times with TBS/T (0.1% Tween20, Sigma #P5927; 20mMTris-HCL pH 7.6, T7943, Sigma, 150mM NaCl, Sigma) then stained with Ponceau S (PoncS) (0.1%w/v in acetic acid, Sigma, #P7170) for 5 min, destained twice for 5 minutes each with 5% acetic acid (v/v) and then twice for 5 minutes with water. Membranes were probed with the following unconjugated antibodies: IkBβ (Santa Cruz, #sc-945), pERK (Cell Signaling Technologies, Danvers, MA, #4370), ERK (Cell Signaling Technologies, #9102) and anti α-actin (Sigma, #A5441). The secondary antibody used for all except anti α-actin was peroxidase conjugated AffiniPure goat rabbit IgG at a dilution of 1:100,000 (Jackson ImmunoResearch Laboratories Inc., West Grove, PA, #111-035-144). The α-actin secondary antibody was a Cy-3 conjugated AffiniPure donkey anti-mouse at a dilution of 1:500 (Jackson ImmunoResearch Laboratories Inc. #715-166-150). Bands for p-ERK, ERK and IkBβ were detected using Amersham ECL Prime Western Blotting Detection Reagents (GE Healthcare, #RPN2252). Gels were blocked with 5% non-fat milk in TBS/Tween. Blots were washed 5 times for 5 minutes each in TBS/Tween. All primary antibodies were incubated for 16 hours with shaking at 4°C. The following concentrations of antibody were used: IkBβ and IkBα 1:200; pERK and ERK 1:1000.

### Gel and Blot Imaging

All images were captured using a Syngene G:Box Chemi XX9 image documentation system and GeneSys software. Image capture settings were auto manual, serial images, and saturation detection specifically using the following sub programs: Chemifast (chemiluminescence), fluorescent (chromophore Cy-3; EPI-Green-M illumination filter module), Coomassie Blue (stained gels) and Manual (PoncS stained membranes). Pixel binning 3×3 (1.01MP) was employed with the Chemifast subprogram. Data analysis/densitometry was performed using the GeneTools program (Syngene). Background subtraction used a “rolling disc” method set at 30 pixels. The imaging software integrates the total pixels in representing the protein band. The resulting “volume density” values, termed RawVol, are the base values used throughout the study. The RawVol values of every target protein band from each lane were added together and averaged. For single bands, IkBβ and actin, this is the band itself; for doublets p-ERK and ERK, both bands are summed and averaged, and for the actin, gel and PoncS total lane staining, all the bands detected were summed and averaged. Normalization of each blot by PoncsS or gel staining was done by determining the median of the lane-averaged values, which was used as the reference value. Each blot lane value was divided individually by the reference value and these proportionalities were used to calculate the normalized value of the target proteins, p-ERK, ERK and IkBβ. Un-normalized or PoncS normalization was used nearly exclusively in this study, (see Experimental Design). All the RawVol and PoncS normalized values used are in the S9 Table.

### Statistics and Graphs

Microsoft Excel (Professional Plus 2013 32 bit) was primarily used for data calculations including means, standard deviations (SD), coefficients of variation (CV), Max/min values, and normalization calculations (S5 Figure). Graphs were produced using Excel or Prism Graph Pad 7.05 running under Windows 10. The NESTED procedure (SAS Institute Inc., Cary, NC) was used to estimate variance components associated with pair, gel (nested within pair), and lane (nested within gel and pair). Percent of variance was calculated by comparing each variance component to the total variance.

### Labels

Figure titles and table annotation are labeled throughout the Results and Supplements based on the data analysis method used in sequential order of the calculations applied using the naming conventions as described in S6 Figure.

### Experimental Design

Supplemental Figure (S3 Figure) illustrates the arrangement of control and treatments on the gels and the number of gels and their pairing for Sample Sets 1 and 2 used to obtain the primary RawVol data. Sample Set 1 and 2 density values are from WB of samples from cell lysates from cells treated with LPS to induce MAPK and NFkB cell signaling pathways or not treated, as controls. MAPK and NFkB were chosen to provide two different responses in RAW cells when exposed to LPS, one which increases, p-ERK, and one that decreases, IkBβ. They differ in that proportion values are either theoretically unbounded or bounded. p-ERK are unbounded as they can be any value equal to or greater than the control. IkBβ values are bounded, as values must be between the control value and zero. Also measured was the total density in each lane from PoncS staining of all the separate bands identifiable on the blot transfer membrane. Gel protein staining prior to blotting and actin density values were also assayed but not used in favor of blot PoncS staining measurements [9, 18, 24-27].

For Sample Set 1 six WB gels were used for analysis with gels loaded with aliquots from either a large pool of control cell lysate or large pool of LPS treated cell lysate. Replicate control and treated aliquots were loaded alternately in the eight lanes not used for molecular weight standards. Gels and transfers were run in pairs as the apparatus can hold 2 gels or 2 transfers at a time. Sample Set 2 used lysates from four separate experiments performed at different times, using independent cultures treated or not treated with LPS which were not pooled. Each experiment was analyzed using two replicate gels with lanes loaded with the same alternating control and LPS pattern for a total of 8 gels producing 8 blots. Therefore there were 2 gels and 2 blot pairs for each of the 4 individual cell culture experiments. In addition to p-ERK and IkBβ, density values for un-phosphorylated ERK (“parental”) protein were determined as was PoncS lane staining, used as the loading control.

All Sample Set 1 samples for the respective control or LPS lanes on all gels were identical and therefore densitometry values should be equivalent, with LPS treated p-ERK values being greater and IkBβ values being less than control values. Theoretically this would result in identical WB density values for all control samples and identical values for all LPS samples. Resulting Max/min ratios of density measurements would be 1, and coefficients of variation would be zero (S4 Figure, S5 Figure). Actual data were only expected to approach these values, of course, but the major study goal was to determine what types of data analysis would result in the closest approach to these ideal values. Sample Set 1 variation should be primarily due to variability caused by the WB analysis process and should demonstrate which analysis process, e.g. work flow, resulted in the best combined method to reduce analytical variation and closest approach to the ideal Max/Min and CV values. Sample Set 2 was done to more closely mimic an actual experimental series by using samples produced from experimental replicate cultures plated, treated, and collected independently on different days, followed by WB analysis using analytical replicates for each experiment. Sample Set 2 allows testing as to whether methods based on Sample Set 1 remain useful. Results from Sample Set 2 were also expected to approach these theoretical ideal values, but not reach them, and differ more from the ideals of Max/Min and CV than results for Sample Set 1 due to purposely introducing additional sources of variation.

By binning the blot results differently for calculations there was the opportunity to determine whether different patterns of sample loading were more or less advantageous (S4 Figure A vs. C and B vs. D). The most typical arrangement is all experimental treatments are run on single gels with analytical replication or replicate experiments on additional gels (S4 Figure A Gel1). But when there are too many different treatments for a single gel such as time series data or complex multiple treatment combinations, then experimental treatment analysis requires the use of multiple gels. As multiple gels are necessary it then is possible to use different methods of grouping replicate samples. Likely the most typical approach is control and treated matched sample pairs being placed in side-by-side lanes with additional pairs of lanes used for each successive treatment of interest. When all lanes have been used the additional treatment pairs are run on additional gels. Analytical replicates of the same samples are loaded using the same pattern on yet additional gels (S4 Figure A Gels 1-6). In short hand, lanes are treatments and gels are replicates.

Alternatively all experimental replicates of a particular treatment or treatment pair could be placed on a single gel (S4 Figure C) or set of additional gels. Additional experimental treatments, with all of their experimental or analytical replicates, would be run on an additional gel or sets of gels (S4 Figure C and D Gels 1-6). In short hand, lanes are replicates and gels are treatments. This arrangement might be a means to reduce variability, as is done when all treatments and replicates can be run on a single gel. A potential disadvantage is this arrangement requires all independent experimental samples be available prior to WB analysis. In the more typical arrangement used, lanes as treatments, WB analysis can be done as samples and analytical replicates from each independent experiment become available and early results evaluated prior to additional experiments. However, in either method of sample arrangement results have to be compared across both lanes and gels. Also in either arrangement there is the potential that sets of two gels can or should be handled as pairs as most electrophoretic apparatus accommodate two gels or two blots. Furthermore a straightforward method of analytical replication can be done using the same set of samples on two gels electrophoresed and transferred together in the same apparatus (S4 Figure B and D).

## Results

### Raw Densitometry Values

Figure 1 shows the RawVol density data from Sample Set 1 and Sample Set 2 for p-ERK and IkBβ. Panels A, B, G, and H show both the within-a-gel lane variability and the variability between gels for every individual lane. The greatest difference is between gels whereas lanes on each gel are similar to one another. This results in the large SD for individual lanes when averaging across gels, as shown by the larger SD bars in Fig 1 C, D, I and J (lanes 3-10). Averages across lanes on each gel result in smaller SD bars for each shown by the comparatively smaller bars for each gel in Fig 1 E, F, K, and L (gel 1-6).

**Fig 1.**
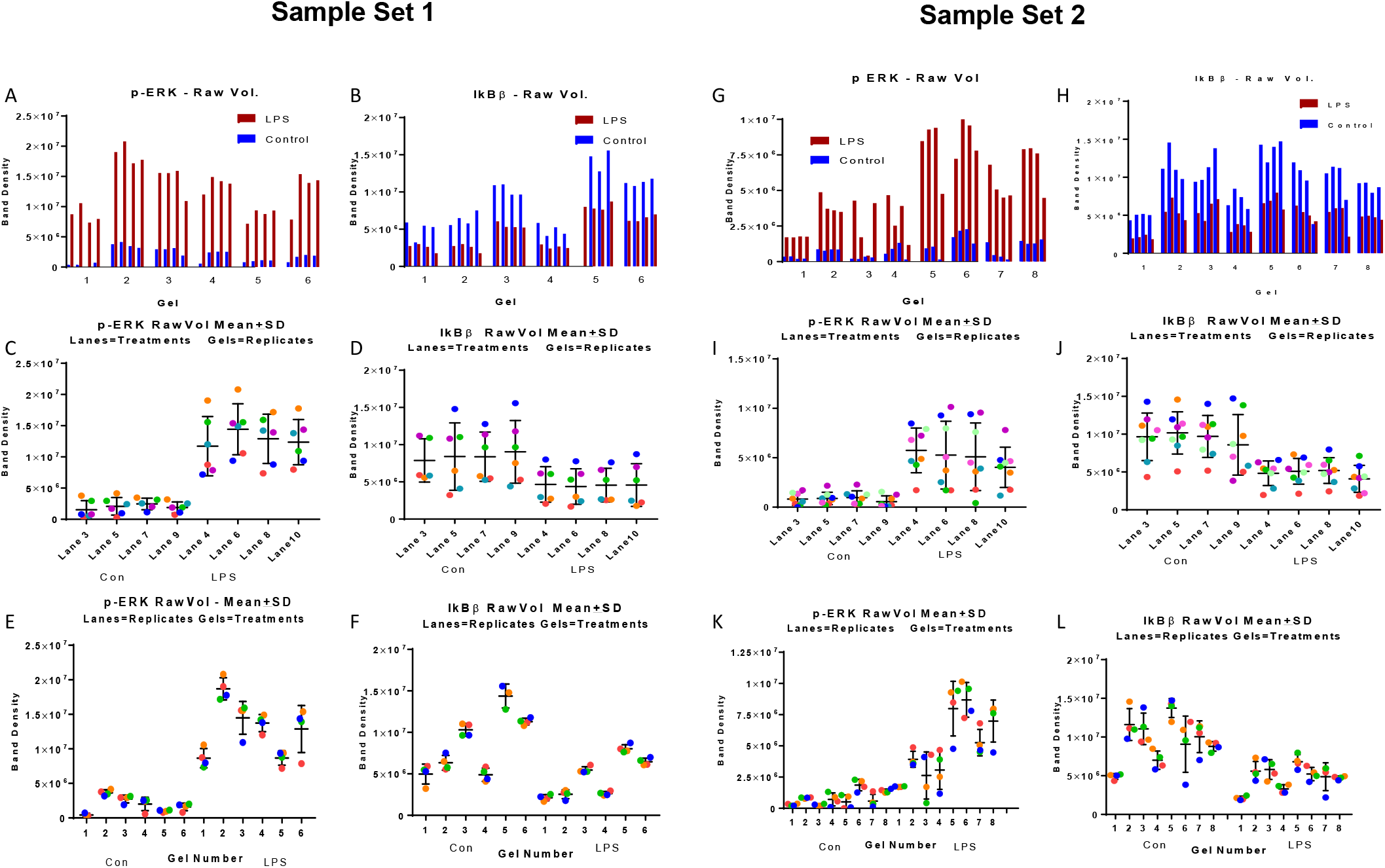
Sample Sets 1 and 2 data without normalization showing individual lane and gel variability. (A, B, G, H) Individual lane density values for all 6 gels for Set 1 (A, B) and Set 2 (G, H) for p-ERK (A, G) and IkBβ (B, H). Values are in the same order as positioned on the gels and resulting blot membranes. Note p-ERK pairs of lane values for control (blue bars) are low and are increased by LPS treatment (red bars); IkBβ pairs of lane values for control (blue) are high and are decreased by LPS treatment (red). Band density is in arbitrary units as rendered by the gel imaging system. (C-L) Graphs group density data by control versus LPS treatment and by using lanes or gels as treatments with replication. Mean and SD values are shown as a vertical line and cross bars respectively for each lane or gel as appropriate. (C, D, I, J). Individual raw data values are grouped with lanes as treatments (3-10) with replicate determinations on gels (1-6) (see S4 Figure A). Dots are individual values from each gel with colors in spectrum order from Gel 1 (red) to Gel 6 (violet) for both control (lanes 3, 5, 7, 9) and LPS treated samples (lanes 4, 6, 8, 10). (E, F, K, L) Individual raw data values are grouped with gels as treatments (gels 1-6) with replicate determinations as lanes (3-10) (see S2 Figure C). Dots are individual values from each lane colored in spectrum order from lanes 3 and 4 (red) to lanes 9 and 10 (blue).

Data variation is also shown by the spread of individual values (color-coded dots) from all 6 gels for that lane (Fig 1 C, D, I, J). However, there is considerable consistency of the response for all the lanes on a particular gel as illustrated by the smaller SD bars for individual gels (Fig 1 E, F, K, L) and the small spread for the individual lane values (colored dots) for most gels, particularly evident in Fig 1 E and F. The mean values are much more consistent when lanes are treatments (Fig 1 C, D, I, J) as despite the large variation between gels, the differences in magnitude for any particular lane relative to other lanes is similar for all gels. For example in Fig 1C the Gel 2 (orange dot) values are the highest values for all four lanes whereas Gel 1 values (red dot) are among the lowest values (cf. Fig1 A and C). This consistency of lane values relative to one another, but considerable variation between gels, results in the mean lane values when averaged across gels being similar but the SD being broad (Fig 1 C). Conversely the mean values are very variable when gels are treatments but the SDs are smaller. The mean treatment values are the average of lanes on a single gel which are similar, but the gels differ considerably. For example Gel 1 and Gel 2 means are very different (Fig 1E) but the lane values on each gel are similar in magnitude to one another, as shown by the tight clustering of individual values. The relative small differences between lanes on different gels is reflected in the relatively smaller SD values and pattern of individual values. For example in Fig 1E Lane 2 values (orange dot) are highest for most gels, and lane 1 values (red dot) being lower, with both varying widely in value on different gels, but are similar to one another on each gel (cf. Fig 1E LPS Gel 1 and 2). Sample Set 2 values show a similar overall pattern, but with even more differences between gels especially for p-ERK. Presumably this reflects the biological variation between independent experiments. Note that pairs of gels are similar to one another while differing from other pairs (cf. Fig 1 G gels 1 and 2 versus 5 and 6).

Analysis of variance of data from Sample Set 1 confirmed both lanes and gel differences contributed to the variance in mean density values, but with 82% attributable to gels-difference and 18% to lane differences (Table 1 RawVol). This statistical result and the patterns in Fig 1 highlights the problem of WB measurement variability when experiments require comparisons across multiple gels; considerable variation occurs even when all the samples being analyzed are biologically identical. In Fig 1 C & D the large SD would preclude discovery of small changes between treatments. In panels E & F erroneous conclusions about time series or treatment effect might be concluded.

**Table 1.** Analysis of Sources of Variance. Sample Set 1 data for four unpaired and one paired set of values. Different normalization methods were tested using nested random effects analysis of variance. Values are the percent of total variation contributed by differences between lanes, gels, and gels grouped in 3 pairs.

| Factor | RawVol | PoncS<br>Norm | Raw Vol<br>Sum Norm | PoncS Sum<br>Norm | Raw Vol<br>PoncS Sum<br>Norm Ratio |
| --- | --- | --- | --- | --- | --- |
| Pair | 0 | 0 | 0 | 0 | 0 |
| Gel | 82.2 | 62.3 | 0 | 0 | 0 |
| Lane | 17.8 | 37.7 | 100 | 100 | 100 |

Comparing ranges of CV values, Sample Set 2 are slightly greater than Sample Set 1 as would be expected with the addition of variation from individual experiments. But RawVol CV values are within a similar range (cf. values in S1 Table and S2 Table); for example, LPS p-ERK lanes as replicates have CVs of 28-41 for Sample Set 1 and 39-67 for Sample Set 2. Therefore the addition of biological variability from independent experiments did not greatly increase the variation observed. Lane variations were greater for IkBβ compared to p-ERK, but IkBβ had slightly less between-gel variations. The biological basis for this difference is likely complex, reflecting the intracellular pool sizes, enzyme kinetics and regulation of the production of p-ERK versus IkBβ, but is beyond the scope of this study. Relative to sample grouping, the preliminary result shows if all experimental treatments and their replicates can be accommodated on a single gel it is advantageous. But cross-gel comparisons are significantly affected by methodological variations. The RawVol data taken together suggest that the primary problem to solve is variation between gels resulting from the WB process.

### Sample Set 1 Normalizations

Starting with the RawVol values a variety of normalization methods were applied to the Sample Set 1 density values. The resulting CV and Max/Min ratios were compared to one another and their convergence to the ideals of 0 and 1, respectively. The results for p-ERK and IkBβ are illustrated in Figure 2 and values are in S1 Table, with more detail in the S 7 Table. Labels on graphs and in Tables are used to describe what calculations have been applied successively to the RawVol density values (S6 Figure). Data were calculated using both lanes as treatments and gels as treatments.

**Fig 2.**
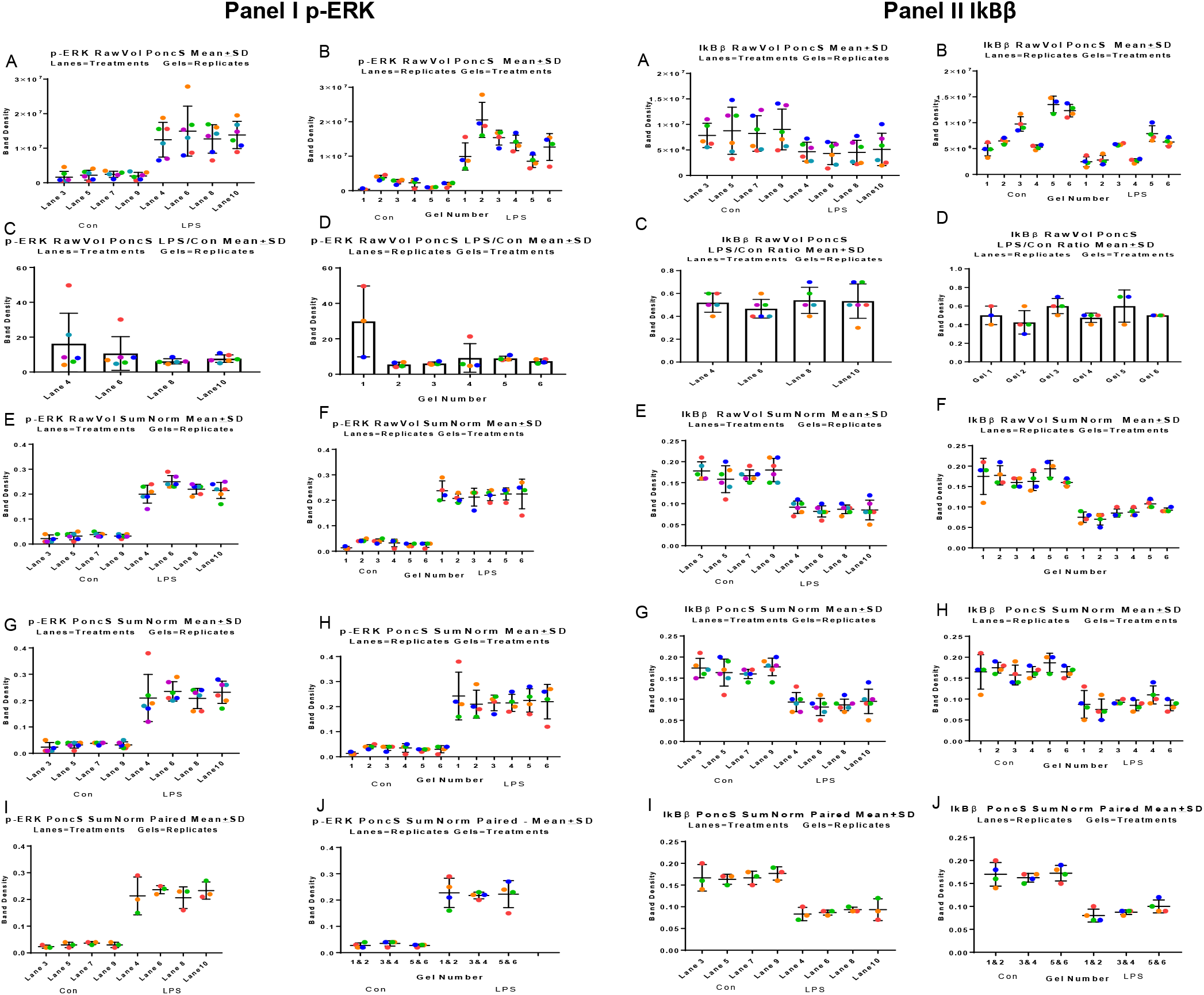
Comparison of multiple normalization methods using Sample Set 1. Means, SD, and individual values, including color coding are as in Fig 1. Mean values closest to equivalency, smallest SD, and least dispersion of individual values are the methods with greatest efficacy. Graph labels describe normalization of raw data (RawVol) with label order of calculations (see S5 Figure and S6 Figure). Initial RawVol starting condition was dropped from overly long titles. Panel I shows p-ERK values and Panel II shows IkBβ values. In each panel (A, C, E, G, I) are for lanes as treatments with gels as replicates and (B, D, F, H, J) are for lanes as replicates with gels as treatments. In the last set of graphs (I, J) in both panels, data were additionally grouped using values for gels in pairs reducing the number of individual gel values from 6 to 3 with the number of lanes remaining 4 (see S4 Figure B, D).

Normalization of target density measurements has been by the density of an independent protein value, such as house-keeping proteins, or more recently total lane protein staining (see Experimental Design). This has been widely viewed as the appropriate means to reduce methodological variability between lanes and remains the most common method applied to WB data. PoncS normalization alone failed to have much effect on the variation of either the p-ERK or IkBβ density values as illustrated by the graphs (cf. Fig 1 C & D with Fig 2 I A & II A and Fig 1 E & F with Fig 2 IB & II B). The RawVol CV and Max/Min values for p-ERK show almost no change (S1 Table). For example, in LPS treated samples Lanes as treatments values for p-ERK CV were 28-41 and 28-48 and Max/min were 1.2 and 1.2, for RawVvol and PoncS respectively. For the respective IkBβ values CV values were 50-63 and 40-62 and Max/Min values 1.1 and 1.2. Another frequent data presentation method is to express the treatment as a proportion of the control response for the matched WB values. For p-ERK this was one of the least effective means of reducing variability for either using lanes or gels as replicates. Max/Min values exceeded 2 and CV were as high as 105 (Fig 2 I C & D, S1 Table p-ERK). This likely reflects that the p-ERK control levels are low, resulting in comparatively weak WB values, which are up to 10 fold lower and more variable than the stimulated response from the LPS-treated samples. Division of the LPS values by the small but variable control values resulted in increased, not decreased variance. In contrast the use of proportional values was effective for IkBβ. The difference likely is IkBβ LPS treated and control values are only approximately 2 fold different resulting in a ratio between two strong WB signal responses. Max/Min ratios were reduced, approaching 1; notably the gels as treatment Max/Min ratio changed from 3.8 to 1.4. CV values were essentially unchanged for gels as treatments or decreased for lanes as treatments (Fig 2 II C & D, S1 and S 7 Table IkBβ).

A normalization method for WB informed by those used for genomic and proteomic arrays was also used [19], termed here Sum Normalization (SumNorm; S5 Figure). This has the advantage of using only the target protein density values themselves and not necessarily requiring the use of protein marker lane normalization. Briefly the sum of the target protein of interest densities are summed across all lanes of the WB, regardless of treatment, and that total used to normalize each lane value on that gel. This method was the best method for reducing variability and was especially effective in reducing the between gel variations as seen by the reduction in SD bars for lanes as treatments (Fig 2 IA vs IE and IIA vs IIE) and the very notable change in the range of mean values for gels as treatments (Fig 2 IB and IF and IIB and IIF). The CV ranges for lanes as treatments for LPS samples are 8-19 for p-ERK and 10-26 for IkBβ, both considerably lower than for the RawVol data (S1 Table). There is not a reduction in CV values for gels as treatments for LPS-treated cells for either p-ERK or IkBβ as these CVs are based on the variation between lanes; as discussed above there is comparatively low variation of lane values within individual gels already. Considering Max/Min values, for lanes as treatments SumNorm does not have much impact; p-ERK value is slightly greater, 1.2 to 1.3 and no change from 1.1 for IkBβ, For gels as treatments SumNorm does impact Max/Min values, bringing them closer to 1, with changes from 2.2 to 1.2 for p-ERK and from 3.8 to 1.5 for IkBβ. Therefore SumNorm reduces the most variable parameter, CV for lanes as treatments or Max/Min for gels as treatments, by about 50% of the RawVol values. All CV are less than 30% and Max/Min values are between 1.1 and 1.5, which are lower than un-normalized data or normalization methods currently used more widely. As current practice is still largely to normalize data by a loading type control the effect of PoncS normalization of lane values was combined with the SumNorm calculation. For p-ERK and IkBβ the result was to increase the SD and broaden the range of CVs (cf. Fig 2 I and II E vs G and F vs H). This likely reflects introducing another source of variation, the PoncS lane sum density values. Max/Min ratio for IkBβ increased slightly or was unaltered, but for p-ERK LPS it beneficially was reduced to 1.1 (S1 Tables). However the CVs and Max/Min values are still closer to the theoretical best values than for un-normalized RawVol, PoncS or Treatment/Control methods.

### Sample Set 1 Data Pairing

The above calculations have treated each gel as a separate independent sample replicate, analogous to six independent experiments. However the WB were actually run as three sets of pairs as the gel electrophoresis and blotting apparatus can accommodate two gels or gels and transfer membranes, a common configuration. Therefore we did calculations based on sets of values being paired as being from gels and blots run at the same time in the same apparatus. This grouping of the six gels was treated as three experiments, each having two analytical replications, and are termed “Paired” in the graphs and tables (Fig 2, S1 Table). Use of paired analytical replicates combined with SumNorm reduces CV for both p-ERK and IkBβ for both lanes as treatments or gels as treatments compared to unpaired values. Paired values of SumNorm without PoncS normalization had lower CVs than paired SumNorm with PoncS normalization. The effect on Max/Min values was more variable with values being the same or slightly higher or lower. An exception is IkBβ where pairing for PoncS and SumNorm gel as normalization treatments reduced Max/Min from 1.5 to 1.2 (S1 Table p-ERK and IkBβ).

### Sample Set 1 Summation

Taken together the major calculation factor reducing CV and Max/Min variation was use of the SumNorm calculation. Furthermore, use of SumNorm resulted in lane differences accounting for all the variance (Table 1) effectively eliminating the confounding influence of samples being on different gels. Use of analytical replicates further reduces variation, as expected. Use of lane density normalization, PoncS in this study, can slightly increase the variability. However, the CV and Max/Min values for Unpaired SumNorm, PoncsSumNorm, Paired SumNorm, and Paired PoncsSumNorm are on the whole fairly similar to one another and more similar, and lower, than the RawVol, PoncS only or LPS/Control normalization methods. The values for both parameters remain greater than 5-10 % range, a rule of thumb value desirable for biological assays, or the ideal values of 1 and 0. But the results suggest that using these alternate normalization methods would allow smaller induced changes in target protein levels to be reliably detected. As current practice is still largely to normalize data by a loading type control the effect of PoncS normalization of lane values was combined with the SumNorm calculation. For p-ERK and IkKβ the result was to increase the SD and broaden the range of CVs. This likely reflects introducing another source of variation, the lane sum density values. Max/Min ratio for IkBβ increased slightly or was unaltered, but for p-ERK it beneficially was reduced to 1.1. However the CVs and Max/Min values are still closer to the theoretical best values than for un-normalized RawVol, PoncS or Treatment/Control methods.

### Sample Set 2 Normalizations

The same set of normalization methods was applied to the RawVol data of Sample Set 2 to determine if similar improvements in reducing variability would be obtained despite samples coming from multiple independent experimental replicates. The Sample Set 2 gels were probed for p-ERK and IkBβ as well as ERK, the un-phosphorylated “parental” protein. This was done as phosphoprotein data are also commonly presented as a ratio of the two forms, in this case p-ERK/ERK values [1, 10]; although it has been suggested this can be problematic [28]. Figure 3 shows graphs for p-ERK, p-ERK/ERK and IkBβ. Primarily the lanes as treatment data are shown, but the last row of graphs are results for paired gels as treatments. CV and Max/Min values for both lanes and gels as treatments for p-ERK, p-ERK/ERK and IkBβ are summarized in the S2 Table.

**Fig 3.**
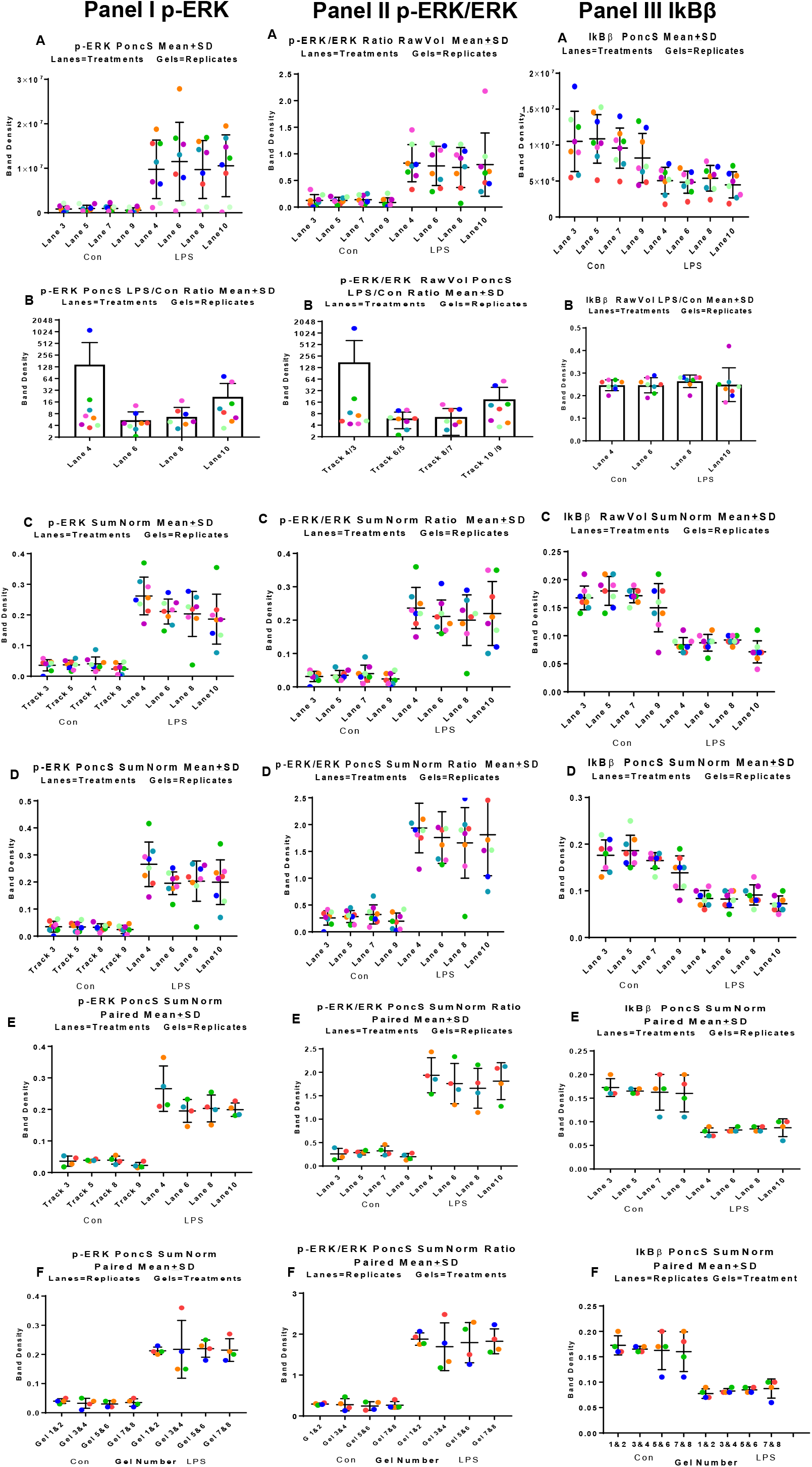
Comparison of multiple normalization methods using Sample Set 2. Means, standard deviations, and individual values, including color coding, are as in Fig 1. Graph labels describe normalization of raw data (RawVol) with labels and order of applied calculations (see S5 Figure and S6). Panel I shows p-ERK values, Panel II shows p-ERK/ERK values and Panel III shows IkBβ values. (A, B, C, D, E) lanes as treatments with gels as replicates and (F) lanes as replicates with gels as treatments. II A p-ERK/ERK show RawVol as PoncS normalization is mathematically equivalent; the PoncS values are factored out in the ratio division. Lanes as treatments are emphasized for brevity as overall pattern of results for gels as treatments was similar for Sample Set 1 (Fig 2) and additional data provided in Fig 4 and S2, S8 Tables.

PoncS normalization alone was not sufficient to reduce CV variability and only slightly reduced Max/Min values for p-ERK, p-ERK/ERK or IkBβ (cf. Fig 3 and Fig 1). Treatment/Control ratio, as for Sample Set 1, was not effective for the p-ERK or pERK/ERK data, increasing both CV and Max/Min values. However, it was still effective in reducing both parameters for the IkBβ data (Fig 3 III B, S2 Table). The pERK/ERK ratio did improve the RawVol Max/Min value compared to p-ERK alone, but resulted in slightly increased CV values (S2 Table). Use of SumNorm with or without PoncS normalization decreased CV values for p-ERK, pERK/ERK and IkBβ compared to RawVol, PoncsS or Treatment/Control. For lanes as treatments Max/Min values were either unchanged or increased slightly. For gels as treatments the effect of SumNorm with or without other normalization on both CV and Max/Min values was substantial, reducing them compared to RawVol values. Notably Max/Min values were as low as 1.1, close to the ideal of 1.0. Pairing also reduced CV across all calculations, but generally not Max/Min values. For Sample Set 2, as for Sample Set 1, paired and unpaired and SumNorm and SumNorm CVs Max/Min values were relatively similar to one another for both gels and lanes as treatments for p-ERK, IkBβ, and p-ERK/ERK. The SumNorm calculation was effective in some cases not only lowering CVs but Max/Min values approached 1 (S2 Table). SumNorm combined with gel lane density normalization has been included in p-ERK/ERK values (S2 Table p-ERK/ERK). This method is similar to using the lane fluorescent intensity of specialized gels for protein staining, but is not equivalent, as staining and imaging occur after, not before, protein transfer to the blot membranes. These values were not reported throughout this study as this staining method would not capture differential lane protein densities, if any, caused during the electro blotting transfer process whereas PoncS staining would be sensitive to this source of change. However, the CV and Max/Min values are identical to those for SumNorm PoncS.

### Sample Set 2 – Combined CV and Max/Min Value Cross Comparison

The study goal is to determine the sample arrangement on WB gels and densitometry calculation methods that together result in the best combined CV and mean value reductions in variability. The results above demonstrate that reductions in CV and of Max/Min values from the RawVol values toward the ideal values of 0 and 1 do not necessarily correlate with successively applied normalization methods; sometimes a further reduction in CV values resulted in a higher Max/Min ratio. Therefore the most useful combination of both needs consideration. To facilitate this Figure 4 summarizes all of the LPS-treated samples for Sample Set 2 data in graphical format to allow ready comparison of CV and Max/Min patterns and combinations. Those chosen for illustration are a subset of all the analyses (S8 Table) and differ slightly between p-ERK and IkBβ based on best combinations of CV and Max/Min values. Values in the p-ERK and IkBβ graphs labeled as ratio are p-ERK/ERK, both LPS treated, and IkBβ/IkBβ for LPS treated/untreated samples, as there is no separate IkBβ form. Non ratio values are results using p-ERK and IkBβ densitometry values from LPS-treated samples. As for the previous CV graphs colored dots are the individual CV values for each of 4 lanes (4,6,8,10) calculated across 8 gels or 4 paired gels, for lanes as treatments (Panel I) or the individual CV values for each of 8 gels (1-8) calculated for 4 lanes, or 4 paired gels, for gels as treatments (Panel II). CV graphs use the same axis range in both panels, as appropriate, for p-ERK or IkBβ, respectively. On Max/Min graphs the ideal value of 1, indicating all densitometry values were the same, is indicated by a red line. This representation also allows the effects of sample arrangement and normalization methods to be readily compared to one another and with starting RawVol values, e.g. the notable change in p-ERK SumNorm CV values for lanes as treatments with or without pairing (Fig. 4 I A and B). Also evident are the consistently large CV ranges for gels as treatments compared to lanes as treatments (Fig. 4 I A and B vs. II A and B) for p-ERK and the consistency of reduced IkBβ CV range and/or values for paired gels as treatments (Fig 4 I F vs. I E and II F vs. II E).

**Fig 4.**
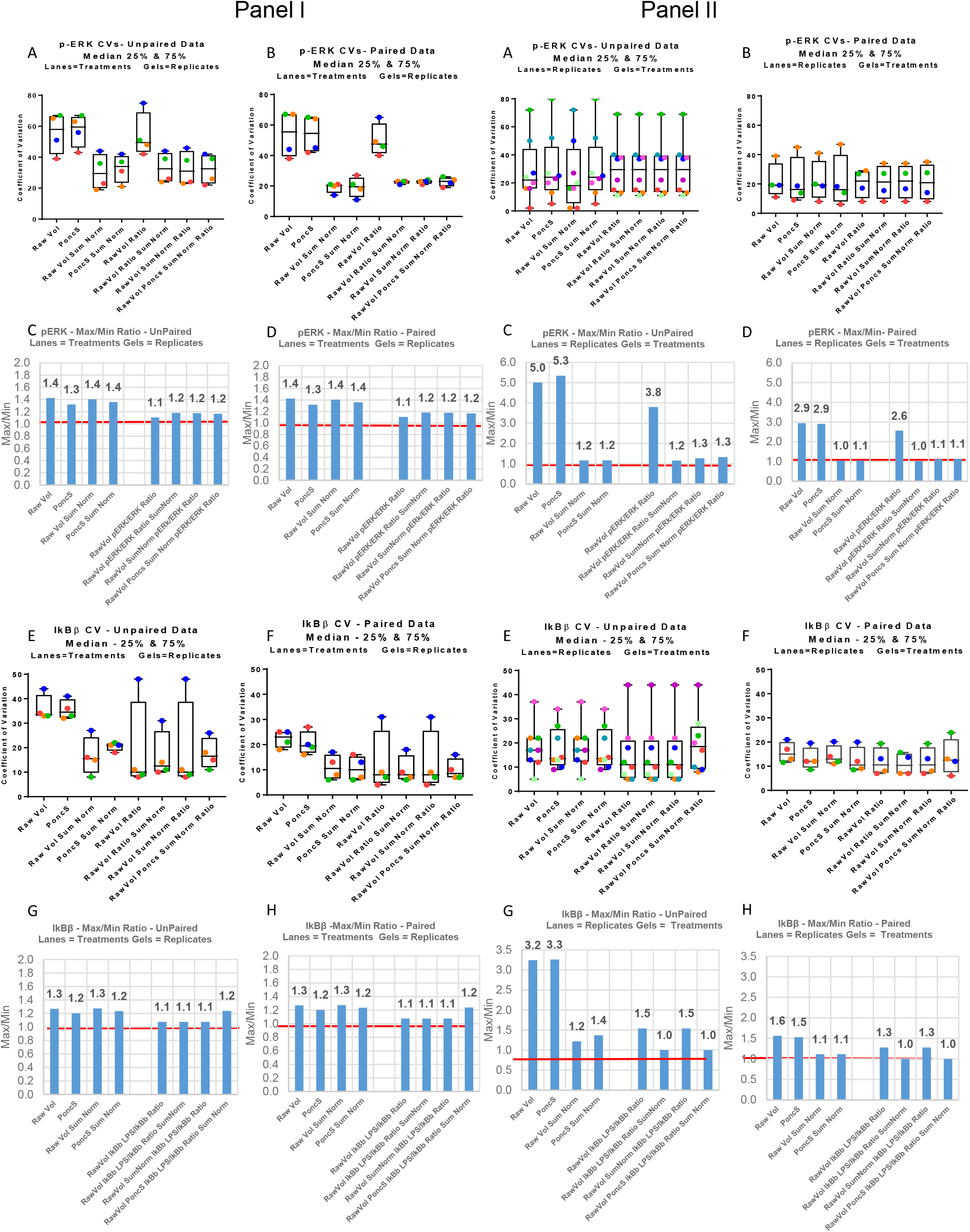
Summary of Sample Set 2 Coefficients of Variation and Max/Min ratios allowing cross comparisons. Summary shows multiple combinations of methods of normalization, pairing, and lanes or gels as treatments in side by side comparison for LPS treated samples. A combination of both the smallest CV range, least individual value spread, and a Max/Min ratio of near or equal to 1 indicates methods most effective in controlling WB methodological variations. CV and Max/Min graphs for the same set of data are positioned vertically together; example A and C and B and D. Panel I (A-H) are data grouped for lanes as treatments with gels as replicates. Panel II (A-H) are data grouped for gels as treatments with lanes as replicates. Panels I and II (A, C, E, G) are unpaired data. Panel I and II (B, D, F, H) are paired data. CV values are shown as box plots with the 25-75th percentiles, max and min whiskers and median value. Mean CV values are nearly the same in most cases. Panel I (A, B, E, F) dots are lane 4, 6, 8, 10 CV values in spectrum order, Lane 4 (red) to Lane 10 (blue). Panel II (A, B, E, F) dots are gel (1-8) CV values in spectrum order, Gel 1 (red) to Gel 8 (violet). Pairing gels results in no change in lane number (Panel I) but reduces gels from 8 to 4 (Panel II). Max/Min Ratio bar charts include the numerical value above bar. Red line is at 1; the value if no difference existed for the mean density value between lanes (Panel I) or gels (Panel II). For each graph normalization description with “ratio”, right side of each figure, indicates p-ERK/ERK or IkBβ-LPS/IkBβ-control. p-ERK and IkBβ values, not used in a ratio are on the left of each figure. LPS treatments only are shown for brevity having the greater absolute value magnitude and dispersion; untreated control values are in S1, S2, S8 Tables.

The lowest Max/Min p-ERK ratios are 1.0 or 1.1, occurring for gels as treatments calculated from SumNorm normalized density values (Fig. 4 II C & II D). Notably this is only true after SumNorm normalization as gels as treatments without this calculation show some of the highest Max/Min values, up to 5.3. However, gels as treatment p-ERK CV values (Fig. 4 II A & B) are constantly large and mostly larger than any values for lanes as treatment for any method of normalization. Therefore, despite the low Max/Min values, due to the large CV ranges, the gels as treatment organization of samples does not provide the best combined CV and Max/Min, regardless of normalization methods. For p-ERK gels as treatments the smallest CV ranges are for p-ERK/ERK paired data with values clustered tightly around 20% (Fig. 4 IB). The p-ERK paired values have individually lower gel CV values and slightly lower median values, for example RawVol SumNorm and PoncS SumNom, but the range is greater (Fig. 4 IB). Max/Min ratios for p-ERK/ERK ratios (1.1-1.2) are consistently lower than p-ERK alone (1.3-1.4), for both paired and unpaired data. The lowest combination of values is for paired p-ERK/ERK RawVol Ratio SumNorm with a Max/Min of 1.2 and very tight CV range with a 20% median (Fig. 4 IB and ID). Nearly as good are the paired p-ERK/ERK values for RAW SumNorm Ratio, where p-ERK and ERK were normalized prior to calculating the p-ERK/ERK ratio. If lane density normalization is required, then the paired PoncS SumNorm with CV of 19-24 and Max/Min of 1.2 is nearly as good (S2 and S8 Tables). Importantly, these results support the idea that experimental treatments with greater than 20% response changes in p-ERK should be able to be discerned.

The smallest and most consistent CV ranges for IkBβ are for gels as treatments paired data, but with little difference whether for IkBβ, alone or as a ratio, with median values of approximately 10% and a range from below 5% to approximately 20% (Fig. 4 II F). Lane as treatment paired CVs are less consistent (Fig. 4 I F), then gels as treatments, but with most SumNorm values for ratio or non-ratio CV median and ranges being comparable to gels as treatments (cf. Fig 4 I F and Fig 4 II F). For most unpaired data, lanes or gels as treatments, CV ranges are greater, sometimes considerably, and with much less consistency, some outliers are in the 40-50% range despite normalization. These largest CV ranges for some normalization methods, for both paired and unpaired, are due to the value from the fourth independent experiment (Fig 4 I E & F blue symbol). A surprising exception is unpaired PoncS SumNorm having the lowest range of CVs values (Fig 4 I E) of the entire IkBβ calculation data set. These two observations suggest that using the IkBβ ratio introduces additional variably, but that either pairing or averaging more values, as for gels as treatments, can reduce the effect of the outlier, in this case the 4^th^ replicate experiment. Max/Min values are mostly 1.3 or lower, many being 1.1, and some 1.0 which indicated no difference between samples. Notable are the gels as treatments RawVol or PoncS, which have the highest values, reflecting gel to gel differences, and this is not compensated by PoncS alone (Fig 4 II G). Application of the SumNorm calculation has the most effect on gels as treatments, bringing Max/Min values closer to 1, again reflecting the between-gel variability that occurs with WB gel replicates. For lanes as treatments, due to the CV variability of paired and unpaired, even with SumNorm, only select unpaired data sets of CV and Max/Min values are optimal choices, particularly the following: unpaired RAW Vol SumNorm (Fig 4 I E&G), paired RawVol Ratio SumNorm and RawVol PoncS SumNorm Ratio (Fig 4 I F&H). Gels as treatments combined CV and Max/Min ratio show good values for paired RawVol SumNorm and RawVol PoncS SumNorm Ratio (Fig 4 II F&H). Considering that lanes as treatments sample arrangement is methodologically simpler than gels as treatments and CV medians and ranges are similar for some normalization calculations, using gels as treatments, despite Max/Min of 1.0 may not be sufficiently advantageous. On that basis the more straight forward and minimal data calculations for paired or unpaired PoncS SumNorm with CV of 10-20% and Max/Min 1.2 may suffice for most studies. As for p-ERK use of SumNorm, pairing, and a ratio of values gave the best result, with considerable improvements of CV values in particular over simple lane protein density normalization.

## Discussion

This study is, to the best of our knowledge, one of the few examining WB performance by explicitly applying the method commonly used to test analytical instrumentation. Uniquely, it combines this approach with evaluation of different sample loading methods and systematic examination of a series of normalization calculations to identify the best combined method for reducing WB analytical (technical) methodological variability. Instrument testing is done with differing goals. One is to define the overall precision and accuracy of the instrument, treating the complex internal steps as a black box, reporting the overall response. The other is to determine the sources of variation for each component and how each step can be improved. Most studies of WB technique to date have focused on the second approach, by making suggestions or collecting data, with the goal of improving individual steps. In this study the first approach was taken, evaluating the overall result of the process, its variability and whether “inputs”, sample arrangement, and/or post-output data processing could reduce this variably. The results demonstrate that considerable control of raw data variability is possible by using appropriate data analysis methods combined with analytical as well as experimental replications. Overall the best approach was to load experimental treatments in successive lanes (lanes as treatments) and use an additional paired gel, processed simultaneously, for analytical replication, for each set of samples. Additional gels pairs are used for sample types not accommodated on a single gel and/or for analysis of additional independent experimental replicates. This physical arrangement is straightforward, and likely already the most commonly used method to generate WB data. Subsequent WB density analysis applied both typical averaging of analytical replicates with gel sum normalization as described by Degasperi et al [19]. This approach was effective in controlling variation for cell signaling responses which were increasing and decreasing and for which control and treatment responses differed in the absolute and relative WB density magnitude. The value of including lane protein density values for normalization was less clear. Purposely multiple methods of data grouping and calculation have been presented to allow others to examine the results and draw conclusions about how best to handle WB densitometry for other target proteins. The Supplementary Material includes the raw densitometry data for PoncS normalized and un-normalized data which can be used for additional calculations to explore other combinations of data grouping and analysis.

The most notable characteristic of the data was the large variations between replicate gels. Although it was expected there would be more gel-to-gel variation than lane-to-lane variation within a single gel, the differences in the RawVol data are striking, especially for the identical samples of Sample Set 1. As experiments were performed by a single individual and the apparatus used for replicate gels in these experiments is the same the results support the inherent complexity of the WB process as an important factor in obtaining highly reproducible density measurements. This result is not surprising based on reviews targeting gel-to-gel variation as an important and problematic source of variation [[15, 16, 19, 29], as well as data variations reported in other studies [1, 10, 18, 20, 21, 25, 30]]. Surprisingly, there seems to be very few reports explicitly targeted towards directly measuring and studying between-gel and between-lane variability of WB. Even more so considering there is a modest literature to examine this type of analytical variability for two-dimensional electrophoresis. In these studies, using direct protein staining, to define CVs, ranges are 5 to 40% [[20, 22, 30] and references therein]. For one-dimensional gel electrophoresis and WB there seem to be few studies to directly estimate variance. Of particular relevance is a study by Deng et al [4] which examined variation in density values between lanes, gels and analysis on successive days of 7 proteins, molecular weight standards, detected by fluorescence. Even without the added complexities of WB they report the detector, dye incubation temperature, agitation during staining and dye batch all contributed to the densitometry measurement and that the variance was different for each protein. In examining lane-to-lane and gel-to-gel variance done under the same conditions for these parameters, both contributed to total variance, but the proportion was different for each protein. Total variance from all factors resulted in CV from 3-13% with lane and gel variability still contributing differentially to the variance for each protein. In a related study CVs ranged from 3 to 38% [23]. In both studies specific steps in the gel staining process were identified as to their contribution to variance, which if controlled could be reduced to 1-3% in some cases for some proteins. In another study using fluorescent protein detection, although statistical parameters were not calculated, data presentation allowing inspection of values for individual gels for multiprotein standard curves shows OD values between analytical replicate gels varying widely, 1.5 to 3 fold [10]. Other studies, while not reporting observed gel differences have described methods to collect all the samples needed for comparison from multiple gels on to the same blot, with the stated purpose being to reduce this source of analytical variability and to increase analytical throughput [9, 17, 29, 30]. These methods were effective but require careful physical cutting of gels and reassembly. As stated in the Introduction, models of error sources and calculation methods to reduce variance have been reported [22, 30], but have not been widely used, perhaps because of computational complexity. An advantage of the approach described here is that it is very similar to normalization methods already widely used.

Results from this study and others suggest that lane normalization with independent protein markers for lane or gel normalization may need to be reconsidered. Most of studies addressing lane normalization have focused on issues with “housekeeping” proteins versus other measurements, but not the overall impact on variation between lanes. The rationale of a loading control to account for differences in sample amounts between lanes seems weak. Total protein measurements of each lane sample should be quite accurate. Modern displacement pipets when used properly have very high precisions and cannot account for the difference in RawVol seen in the Sample Set 1 results (Fig 1). Especially note the differences between pairs of cells such as Gel 1 and Gel 2 in Figure 1 A. Further our experience suggests that even loading 25% of the total protein of a reference sample failed to alter either actin or PoncS lane densities in a linear fashion (data not shown). Therefore variations in loading control are likely only apparent for very large changes in loading and not sufficiently sensitive to correct smaller lane-to-lane variations that could affect comparative target protein measurements. The other reason for use of an invariant protein is to normalize changes caused by the WB steps such as membrane transfer and detection. In this study use of the PoncS lane density values alone did not decrease CVs very much and when combined with SumNorm in most cases increased the CV values (cf. Fig 1 B and Fig 2 IB). The use of a reference sample, of either known concentration or as a baseline to correct difference between gels/membranes, may also need reconsideration. The results showed that although variation between lanes was less than between gels it was almost 18% of the total variation in raw data. Stated differently, due to lane-to-lane variability any particular lane might be the highest or lowest even for successive analytical replicate gels, causing the internal standard to become a shifting baseline. Taken together, although the presence of loading controls is visually appealing on representative blot images, it may not in fact be good evidence of accomplishing their stated purpose of showing methodological consistency, especially when less than multifold changes in the target proteins are of interest. This possibility should be given further consideration and investigated by others.

Due to its continued widespread use, studies to improve WB methodology are still needed. One approach is to change and improve the physical apparatus used to conduct WB. Technical changes that have already occurred for gel electrophoresis and WB include availability of commercial pre-cast gels and buffer stocks, more rapid semi-dry blotters, and gels compounded so all proteins fluoresce. Surprisingly there seem to be no reports making measurements of electric fields within gel electrophoresis boxes either during the gel separation step or the blot transfer step. Differences in field strengths (fluence variations) during either of these processes seem a possible mechanism that could cause lane or gel protein densities to differ. Likewise differential surface properties of blot membranes or gel characteristics may be less uniform than assumed. Prior to conducting these studies we accepted the common assumptions that electrodynamics of solutions allow only small variations within electrophoresis apparatus and uniformity of commercially produced gels and membranes seemed warranted. But these are two areas that investigation might lead to improvements in WB performance. Capillary electrophoresis is a significant redesign of the physical technology to separate and identify proteins that still rely on electrophoretic separation and antibody detection. Commercial applications also couple this physical change with internal markers, novel fixation of proteins and software with normalization algorithms. Published evaluations of these technologies used typical instrument testing approaches to provide estimates of accuracy and precision for these instruments and suggest that there are reductions in variability due to analytical methodology [5, 31]. A limitation is these instruments are comparatively expensive resulting in high entry costs for switching to this technology compared to widely used tank based systems with or without semi-dry transfer. Also the fixed cartridge formats are better suited to studies with simultaneous analysis of larger number of samples. This is also true for antibody capture arrays. Even custom arrays may not have all the targets of interest due to a lack of antibodies that function well and or there can be sensitivity limitations for particular analytes compared to WB. Smaller labs, smaller studies, or those outside of research centers likely will need to continue to use traditional WB methods.

The other approach is to improve data analysis methods. An important information gap in this literature is a lack of multi-gel analysis, with many reports using single gel observations to illustrate methodological problems or improvements. This study benefited from the application of a method derived from analysis of nucleic acid arrays, which makes extensive use of analytical replicates in addition to independent experimental replicates [19]. Data post processing and statistical methods are becoming increasing sophisticated for analysis of various types of arrays, mass spectroscopy, and nucleic acid sequence data. These technologies have both the challenge and advantage of much greater amounts of analytical replication then is typical or even possible for WB based assays. But it may be useful to examine whether there are additional experimental designs and algorithms, beyond those described and referenced here, that could further improve WB analysis.

## Author Contributions

Conceptualization: R. Joel Lowy

Formal Analysis: R. Joel Lowy

Funding Acquisition: R. Joel Lowy

Investigation: Phyllis A. Rees

Methodology: Phyllis A. Rees and R. Joel Lowy

Project Administration: R. Joel Lowy

Visualization: R. Joel Lowy

Writing-Original Draft: R. Joel Lowy and Phyllis A. Rees

Writing - Review and Editing: R. Joel Lowy and Phyllis A. Rees

## Acknowledgements

The Biostatistics Consulting Center, Department of Preventive Medicine and Biostatistics, USU, and in particular Dr. Cara Olsen and Dr. Anwar Ahmed performed the Nested Random Effects Analysis of Variance.

## Data Availability Statement

All relevant data are within the manuscript and the Supporting Information files.

## Supporting Information

**Table S1.** CV and Max/Min Value Summary for p-ERK and IkBβ Sample Set 1. Numerical values for the subset of normalization methods illustrated in Fig 1 and Fig 2. SumNorm Paired values not in graph are included for comparison to SumNorm (unpaired) and PoncS SumNorm Paired results.

| Normalization Method | Lanes as Treatments<br>Gels as Replicates |  |  |  | Gels as Treatments<br>Lanes as Replicates |  |  |  |
| --- | --- | --- | --- | --- | --- | --- | --- | --- |
|  | Control |  | LPS |  | Control |  | LPS |  |
|  | CV | Max/<br>min | CV | Max/<br>min | CV | Max/<br>min | CV | Max/<br>min |
| p-ERK<br>RawVol | 38-95 | 1.6 | 28-41 | 1.2 | 12-57 | 8.4 | 9-26 | 2.2 |
| p-ERK<br>PoncS | 37-103 | 1.5 | 28-48 | 1.2 | 11-53 | 8.9 | 14-40 | 2.4 |
| p-ERK<br>RawVol<br>LPS/Con. |  |  | 20-102 | 2.2 |  |  | 10-81 | 5 |
| p-ERK<br>SumNorm | 16-70 | 1.7 | 8-19 | 1.3 | 12-57 | 3.4 | 9-26 | 1.2 |
| p-ERK<br>PoncS SumNorm | 15-45 | 1.6 | 18-43 | 1.1 | 11-50 | 3.7 | 14-40 | 1.1 |
| p-ERK<br>SumNorm Paired<br>Graph not shown | 10-25 | 1.7 | 9-17 | 1.3 | 21-25 | 1.4 | 9-19 | 1.1 |
| p-ERK<br>PoncS Sum Norm<br>Paired | 9-31 | 1.6 | 7-34 | 1.3 | 20-26 | 1.3 | 6-24 | 1.1 |

IκBβ
| Normalization Method | Lanes as Treatments<br>Gels as Replicates |  |  |  | Gels as Treatments<br>Lanes as Replicates |  |  |  |
| --- | --- | --- | --- | --- | --- | --- | --- | --- |
|  | Control |  | LPS |  | Control |  | LPS |  |
|  | CV | Max/<br>min | CV | Max/<br>min | CV | Max/<br>min | CV | Max/<br>min |
| IκBβ<br>RawVol | 37-54 | 1.2 | 50-63 | 1.1 | 7-24 | 2.9 | 7-21 | 3.8 |
| IκBβ<br>PoncS | 30-53 | 1.2 | 40-62 | 1.2 | 9-26 | 2.8 | 3-39 | 3.2 |
| IκBβ<br>LPS/Con. |  |  | 9-27 | 1.1 |  |  | 4-29 | 1.4 |
| IκBβ<br>SumNorm | 8-20 | 1.2 | 10-26 | 1.1 | 4-24 | 1.2 | 7-21 | 1.5 |
| IκBβ<br>PoncS SumNorm | 6-20 | 1.1 | 16-29 | 1.2 | 9-26 | 1.2 | 3-39 | 1.5 |
| IκBβ<br>SumNorm Paired<br>Graph not shown | 9-15 | 1.1 | 9-14 | 1.1 | 9-32 | 1.2 | 6-10 | 1.4 |
| IκBβ<br>PoncS Sum Norm<br>Paired | 4-15 | 1.10 | 5-26 | 1.2 | 4-35 | 1.1 | 8-17 | 1.2 |

**Table S2.** CV and Max/Min Value Summary for p-ERK, p-ERK/ERK and IkBβ Sample Set 2. Numerical values for the subset of normalization methods illustrated in Fig 1 and Fig 3. Paired p-ERK and IkBβ values not in graphs are included for comparison to SumNorm (unpaired) or PoncS SumNorm Paired results. p-ERK/ERK normalized using electrophoretic gels staining density which was not graphed is also included for comparison to PoncS normalization.

| Normalization Method | Lanes as Treatments<br>Gels as Replicates |  |  |  | Gels as Treatments<br>Lanes as Replicates |  |  |  |
| --- | --- | --- | --- | --- | --- | --- | --- | --- |
|  | Control |  | LPS |  | Control |  | LPS |  |
|  | CV | Max/<br>min | CV | Max/<br>min | CV | Max/<br>min | CV | Max/<br>min |
| p-ERK RawVol | 71-104 | 1.7 | 39-67 | 1.4 | 6-106 | 7.1 | 2-72 | 5.0 |
| p-ERK PoncS | 72-103 | 1.8 | 43-67 | 1.3 | 12-107 | 6.7 | 5-80 | 5.3 |
| p-ERK LPS/Con. |  |  | 43-288 | 28 |  |  | 15-105 | 72 |
| p-ERK SumNorm | 41-74 | 1.1 | 19-44 | 1.4 | 6-106 | 3.1 | 2-72 | 1.2 |
| p-ERK PoncS SumNorm | 44-74 | 1.1 | 21-42 | 1.4 | 12-107 | 3.1 | 5-80 | 1.2 |
| p-ERK SumNorm Paired<br>Graph not shown | 16-40 | 1.7 | 14-21 | 1.4 | 13-53 | 1.3 | 8-20 | 1.1 |
| p-ERK PoncS Sum Norm<br>Paired | 8-45 | 1.7 | 11-27 | 1.4 | 14-50 | 1.4 | 6-18 | 1.1 |

B p-ERK/ERK
| Normalization Method | Lanes as Treatments<br>Gels as Replicates |  |  |  | Gels as Treatments<br>Lanes as Replicates |  |  |  |
| --- | --- | --- | --- | --- | --- | --- | --- | --- |
|  | Control |  | LPS |  | Control |  | LPS |  |
|  | CV | Max/<br>min | CV | Max/<br>min | CV | Max/<br>min | CV | Max/<br>min |
| p-ERK/ERK RawVol | 87-101 | 1.6 | 42-75 | 1.1 | 13-106 | 5.5 | 11-69 | 3.8 |
| p-ERK/ERK PoncS | 49-101 | 1.1 | 42-75 | 1.1 | 13-106 | 5.5 | 11-69 | 3.8 |
| p-ERK/ERK LPS/Con. |  |  | 73-289 | 29 |  |  | 12-188 | 76 |
| p-ERK/ERK SumNorm | 42-75 | 1.7 | 24-44 | 1.2 | 13-106 | 3 | 11-69 | 1.2 |
| p-ERK/ERK PoncS SumNorm | 39-77 | 1.6 | 24-42 | 1.2 | 13-106 | 3 | 11-69 | 1,3 |
| Gel Stain SumNorm<br>Ratio<br>Data not Graphed | 40-77 | 1.7 | 22-42 | 1.2 | 13-106 | 2.7 | 11-69 | 1.3 |
| p-ERK/ERK SumNorm Ratio<br>Paired | 19-42 | 1.7 | 21-23 | 1.2 | 8-55 | 1.2 | 8-34 | 1.0 |
| p-ERK/ERK PoncS Sum Norm<br>Paired | 17-45 | 1.6 | 19-24 | 1.2 | 8-46 | 1.2 | 8-28 | 1.1 |

C IκBβ
| Normalization Method | Lanes as Treatments<br>Gels as Replicates |  |  |  | Gels as Treatments<br>Lanes as Replicates |  |  |  |
| --- | --- | --- | --- | --- | --- | --- | --- | --- |
|  | Control |  | LPS |  | Control |  | LPS |  |
|  | CV | Max/<br>min | CV | Max/<br>min | CV | Max/<br>min | CV | Max/<br>min |
| IκBβ RawVol | 33-47 | 1.2 | 33-34 | 1.3 | 7-40 | 2.8 | 5-22 | 3.2 |
| IκBβ PoncS | 31-42 | 1.3 | 32-41 | 1.2 | 6-37 | 2.8 | 9-27 | 3.3 |
| IκBβ RawVol LPS/Con. |  |  | 9-49 | 1.2 |  |  | 4-44 | 1.6 |
| IκBβ SumNorm | 8-29 | 1.3 | 8-27 | 1.3 | 8-40 | 1.1 | 5-37 | 1.2 |
| IκBβ PoncS SumNorm | 10-25 | 1.3 | 18-21 | 1.2 | 6-37 | 1.2 | 9-34 | 1.4 |
| IκBβ PoncS Sum Norm<br>Paired | 4-22 | 1.3 | 7-16 | 1.2 | 9-24 | 1.1 | 9-20 | 1.1 |
| IκBβ Ratio SumNorm Paired<br>Graph not shown |  |  | 9-18 | 1.1 |  |  | 7-16 | 1.0 |
| IκBβ PoncS Ratio SumNorm Paired<br>Graph not shown |  |  | 7-16 | 1.2 |  |  | 6-23 | 1.0 |

**Figure S3.**
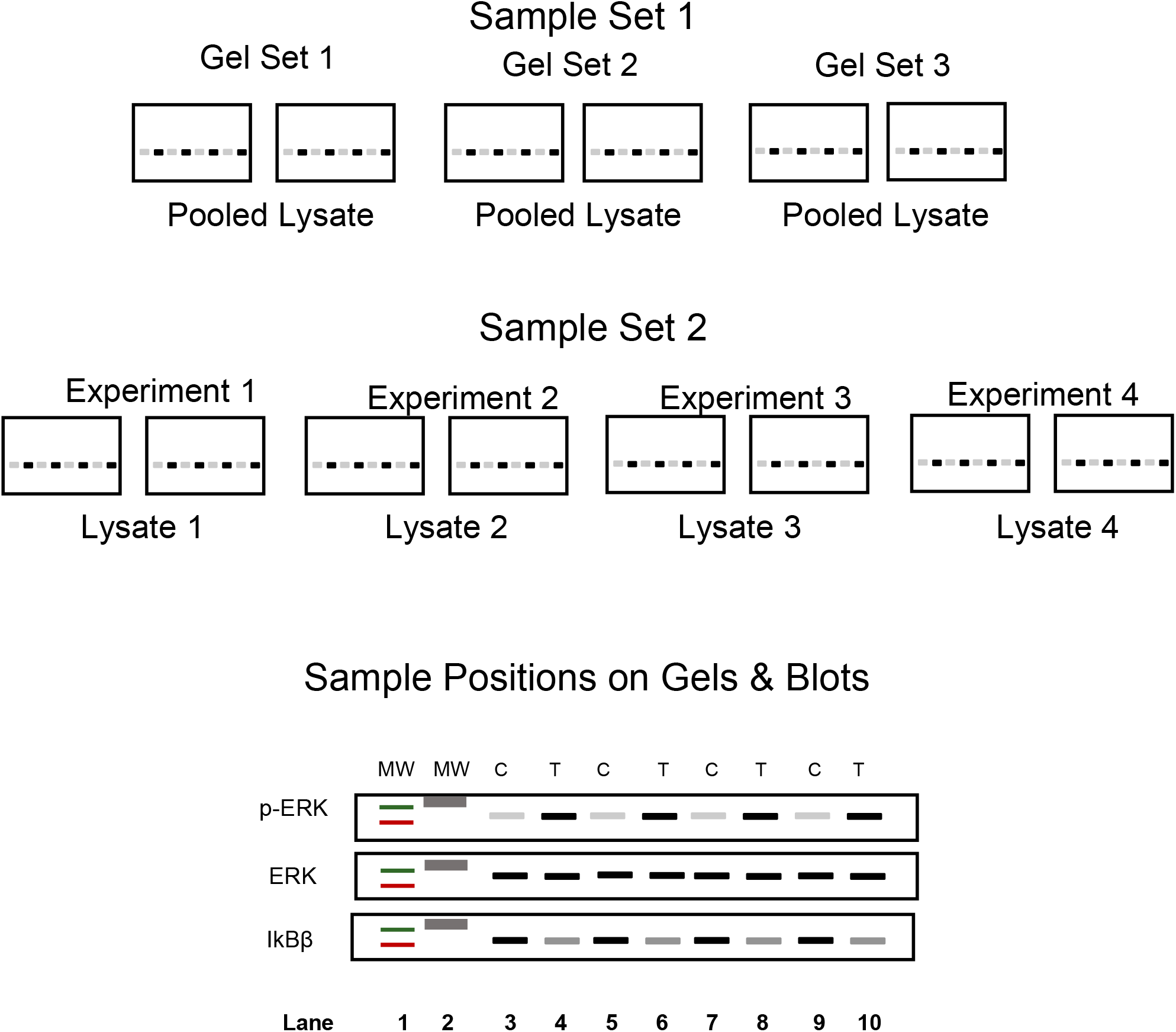
Gel Layouts for Sample Set 1 and Sample Set 2. Graphical representation of arrangements and relationships to one another of LPS treated and untreated samples, lysates on gels, gel numbers and gel pairing. Sample Set 1 pooled lysate was used for all gels. For Sample Set 2 each experiment pooled lysate was kept separate and used for the respective pairs of gels. Lanes 1 and 2 were used for MW markers. Lanes 3, 5, 7 and 9 were loaded with control lysates. Lanes 4, 6, 8 and 10 were loaded with lysates from LPS treated cell cultures.

**Figure S4.**
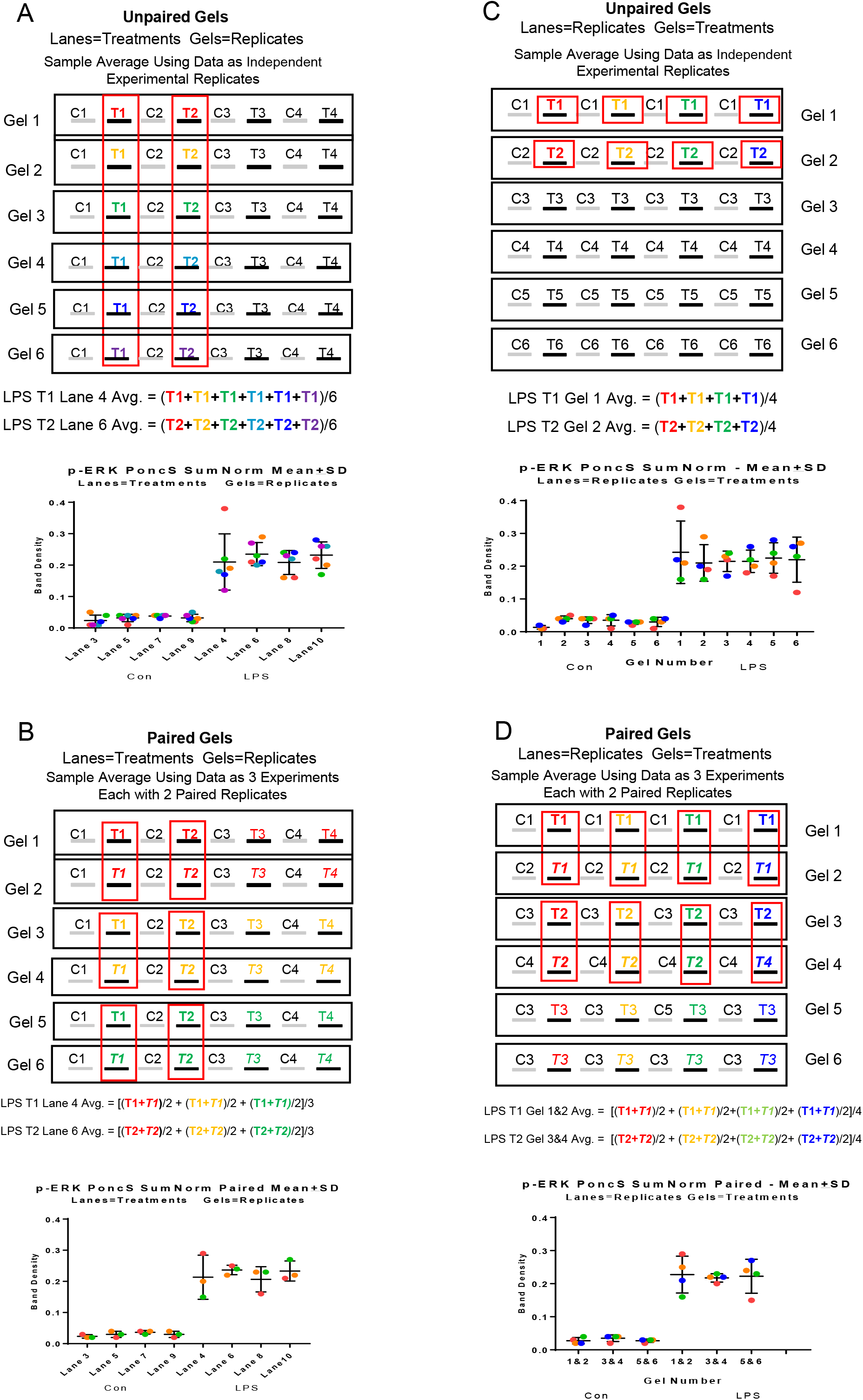
Data Grouping for Lanes as Treatments versus Gels as Treatments. Graphical representation of how individual lane density values are grouped for statistical calculations. Shown are groupings of Unpaired (A, C) and Paired (B, D) for lanes as treatments (A, B) and gels as treatments (C, D). Individual samples are color coded showing how individual density values map from gel/blot to average calculation and as individual values on final graphical representations. For clarity and brevity of illustration only LPS treated samples from six gels and the first two average calculations are displayed. SD and CV calculations use the same groupings.

**Figure S5.**
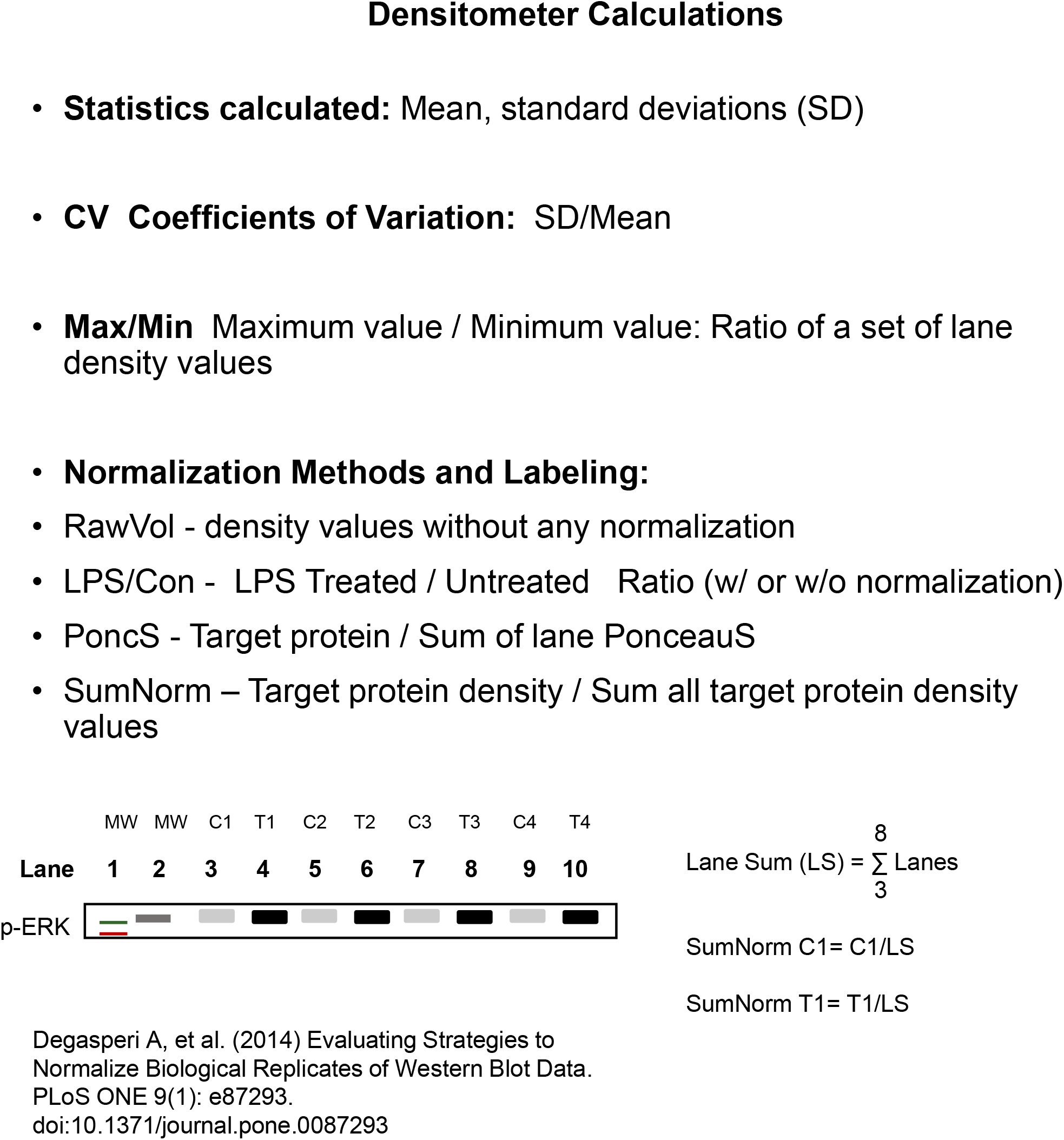
Summary of Densitometry Calculations and Labeling. Synopsis of statistical calculations and the nomenclature that is used for the density values with and without normalization. As an example RawVol values are those from the gel imaging system without any additional calculations applied. The SumNorm calculation formula is shown and illustrated graphically. Also the correspondence between sample number and gel/blot lane number.

**Figure S6.**
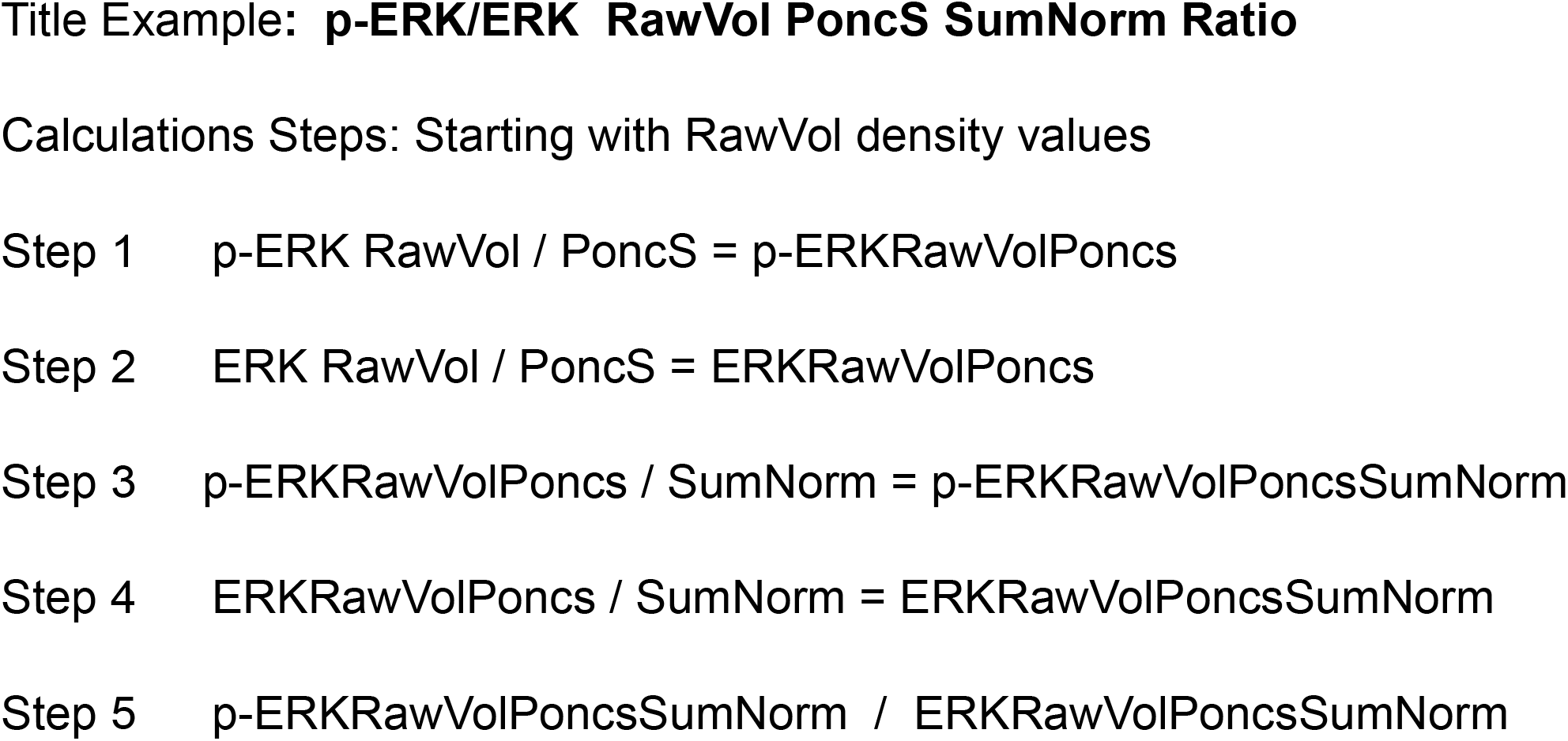
Nomenclature for Graphs and Table Titles. Multiple calculations methods were applied to the raw density values (RawVol) and compared. Titles are descriptive as to which calculations were applied with the order of labels corresponding to the order operations were applied to the raw data. A relatively complex, muti-step normalization is used for illustration, p-ERK/ERK RawVol PoncS SumNorm Ratio. Note that this is not equivalent to p-ERK/ERK RawVol PoncS Ratio SumNorm, which was also used. The difference is whether the SumNorm calculation was applied before or after calculation the ratio of p-ERK to ERK. For some very long titles the leading RawVol was dropped as this is always the initial density value used for analysis for all samples.

**Table S7.**
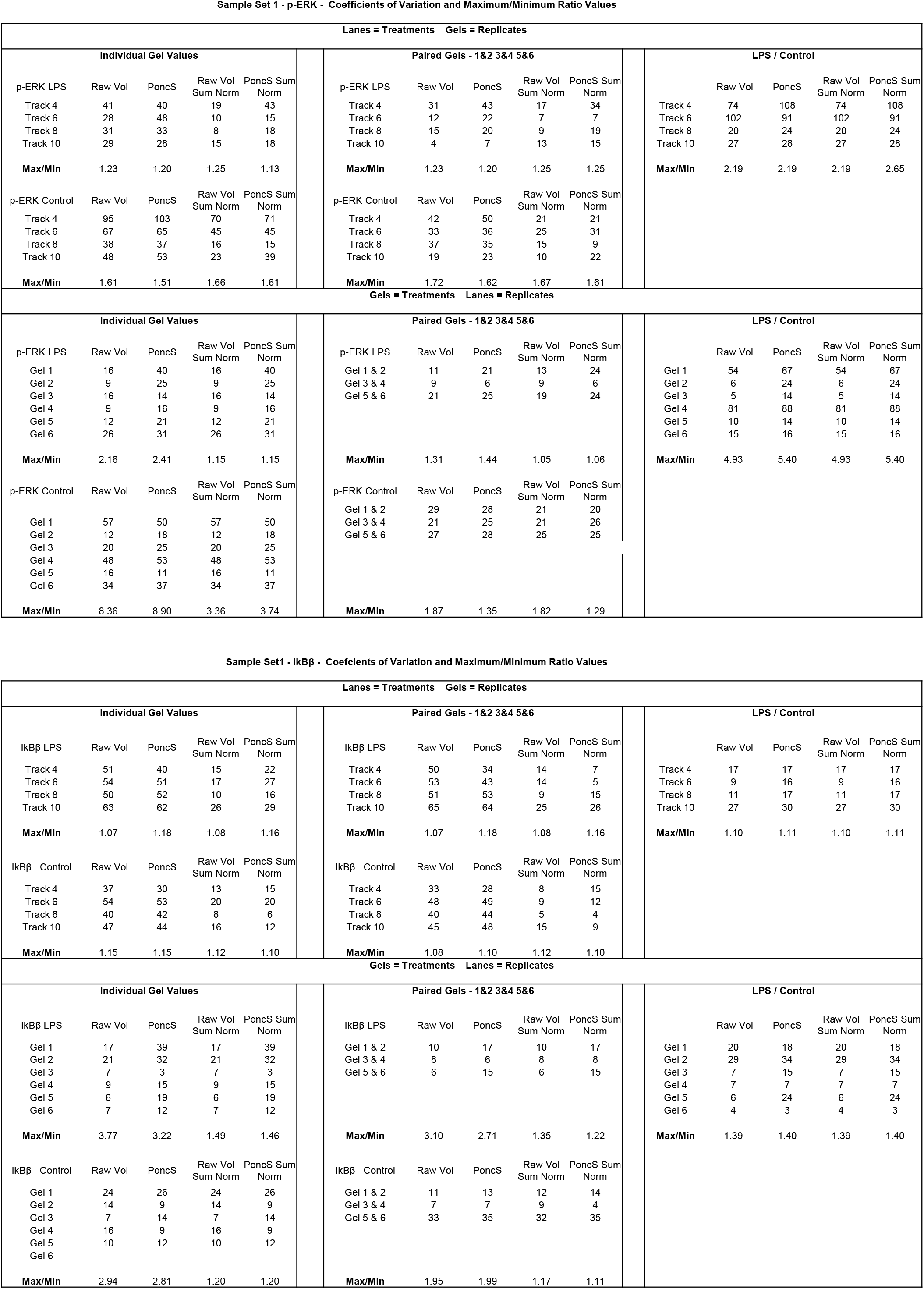
Summary of CV and Max/Min Ratios for All Sample Set 1 Data. Values for all normalization methods, lanes or gels as treatments, Pairing and Ratio combinations are provided. Figures 1-4 and S1 Table are a subset to illustrate the results of different value grouping and normalizations.

**Table S8.**
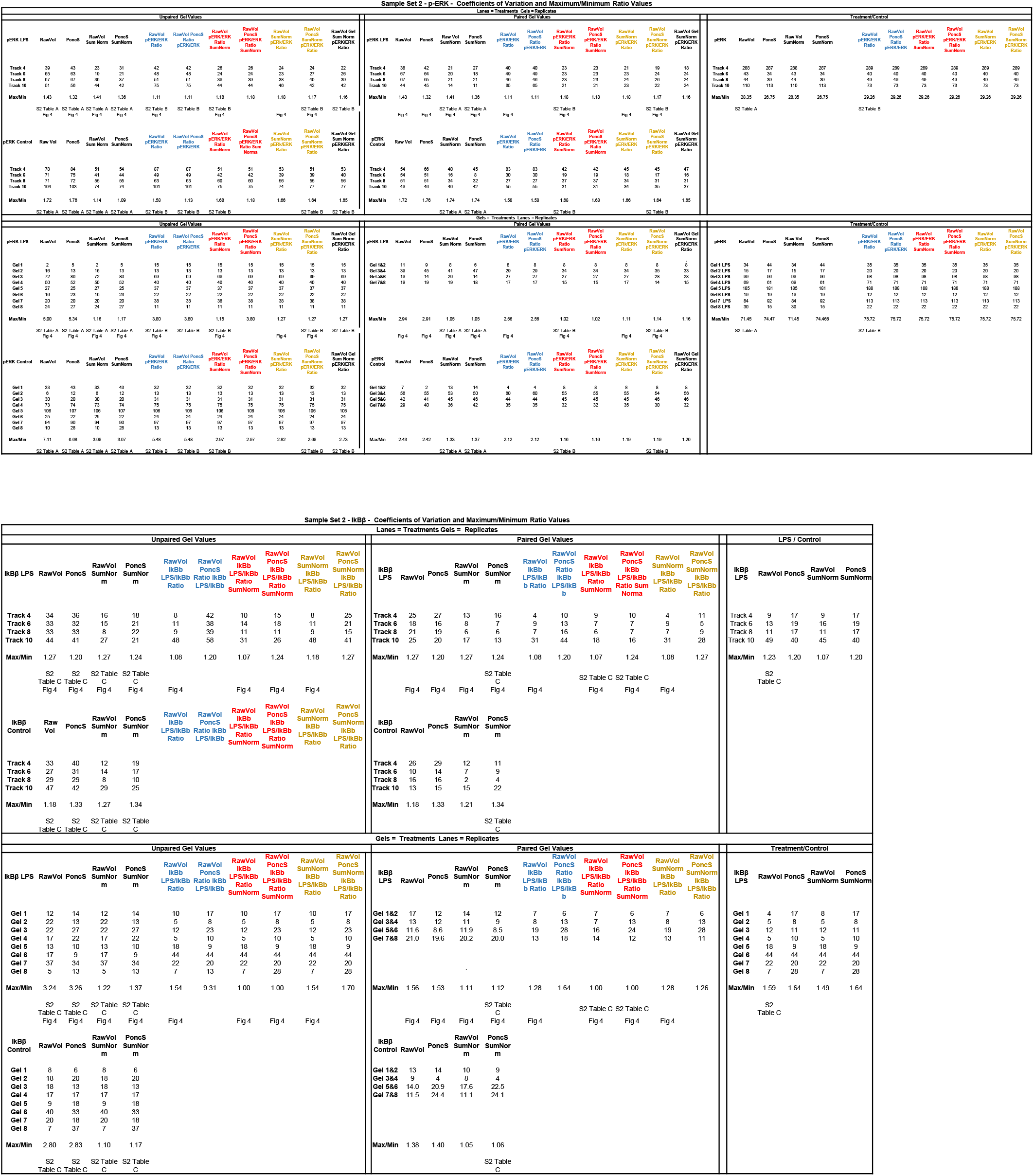
Summary of CV and Max/Min Ratios for All Sample Set 2 Data. Values for all normalization methods and groupings, Lanes or Gels as treatments, Pairing and Ratio combinations used are provided. Figures 1-4 and Supplementary S2 Table are a subset to illustrate the results of different value grouping and normalizations. Values used in the manuscript body figures and S2 Table summaries are annotated. Color Coding for the p-ERK/ERK and IkBβ-LPS/IkBβ-control ratios has been used to clarify which sets of calculations applied are similar. Inspection shows many but not all Blue, Red, and Yellow values are identical or similar, despite the differences in order in which calculations were applied. For some calculations this is due to some ratio effects being canceled out. An example is p-ERK/ERK SumNorm with and without PoncS; as the PoncS was applied to both the p-ERK and ERK values it is factored out in the division calculation and has no effect.

**Table S9.**
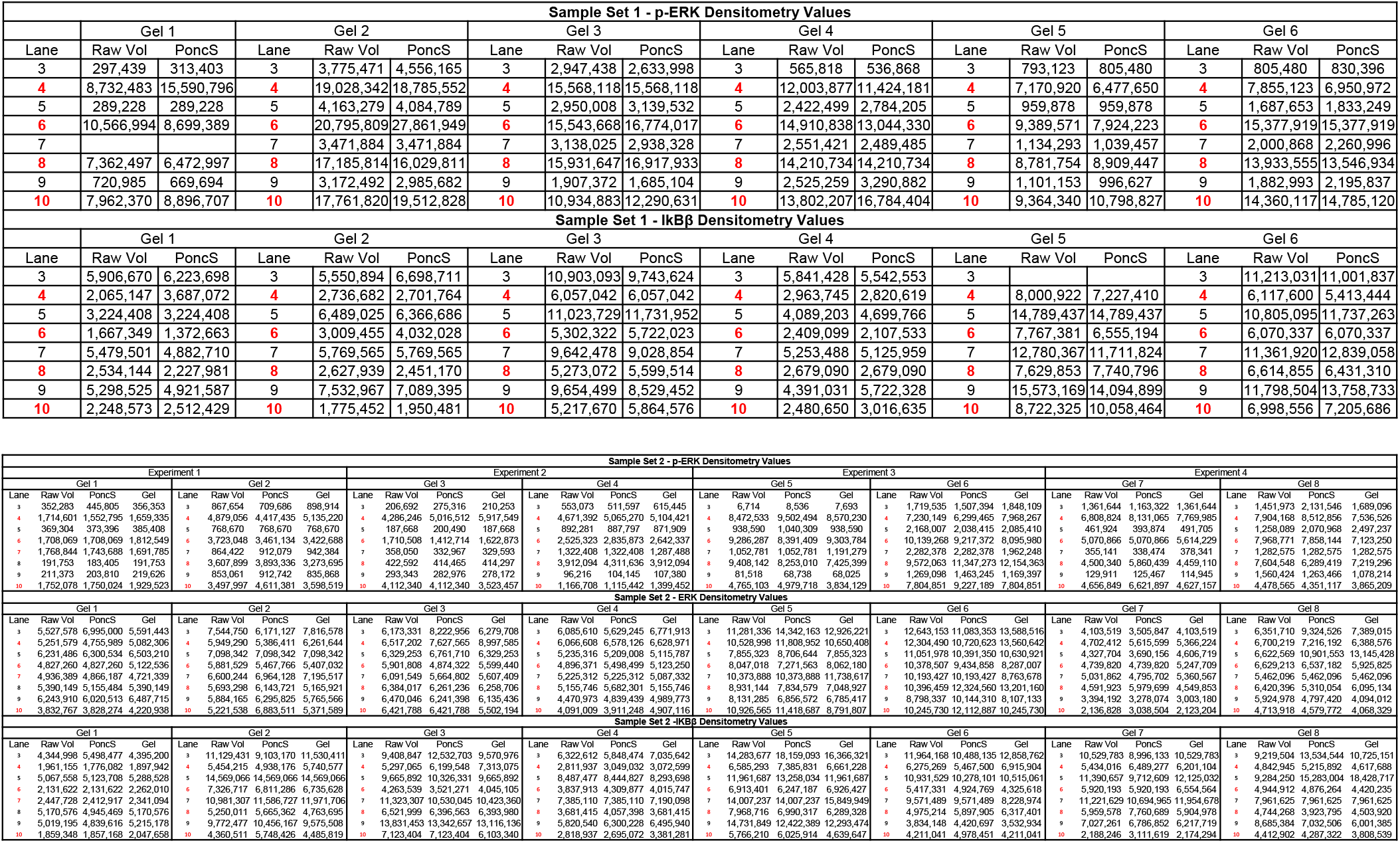
Summary of RawVol Densitometry Values for Sample Sets 1 and 2. All of the initial raw data or raw data normalized with PoncS used in this study for both sample sets are provided. Red lane numbers indicated LPS treatment (4,6,8,10), black (3,5,7,9) are untreated control samples. Lane numbers start with 3 as Lanes 1 and 2 were used for molecular weight markers (see S3 Figure and S5 Figure).

